# Does the posteromedial cortex play a primary role for the capacity for consciousness in rats?

**DOI:** 10.1101/2021.01.22.427747

**Authors:** A. Arena, B.E. Juel, R. Comolatti, S. Thon, J.F. Storm

**Affiliations:** Department of Molecular Medicine, University of Oslo, Oslo, Norway; Center for Sleep and Consciousness, University of Wisconsin, Madison, WI, USA; Department of Biomedical and Clinical Sciences “L. Sacco”, University of Milan, Milan, Italy

## Abstract

It remains unclear how specific cortical regions contribute to the brain’s overall capacity for consciousness. Clarifying this could help distinguish between theories of consciousness. Here, we investigate the association between markers of regionally specific (de)activation and the brain’s overall capacity for consciousness.

We recorded electroencephalographic (EEG) responses to cortical electrical stimulation in 6 rats, and computed Perturbational Complexity Index state-transition (PCI^ST^), which has been extensively validated as an index of the capacity for consciousness in humans. We also estimated the balance between activation and inhibition of specific cortical areas with the ratio between high and low frequency power (HF/LF) from spontaneous EEG activity at each electrode. We repeated these measurements during wakefulness, and under the influence of ketamine anaesthesia at two doses: the minimal dose needed to induce behavioural unresponsiveness and twice this dose.

We found that PCI^ST^ was only slightly reduced from wakefulness to light ketamine anaesthesia, but dropped significantly down with deeper anaesthesia. The high-dose effect was selectively associated with reduced HF/LF ratio in the posteromedial cortex, which strongly correlated with PCI^ST^. Conversely, behavioural unresponsiveness induced by light ketamine anaesthesia, was associated with similar spectral changes in frontal, but not posterior cortical regions.

These findings seem to support the claim that the posteromedial cortex may play a primary role for the capacity for consciousness. Such region-specific associations between cortical activation and the overall capacity for consciousness must be accounted for by theories of consciousness.

## Introduction

It is widely recognized that relatively few parts of the brain are directly involved in specifying our everyday conscious experiences (Koch *et al.*, 2016). Ideally, a theory of consciousness should be able to precisely explain why only these regions contribute. Recently, the roles played by the frontal vs. posterior parts of the neocortex in the consciousness of healthy humans have been discussed (Boly *et al.*, 2017; Odegaard, Knight and Lau, 2017). In this “front vs. back debate”, some have argued that the “evidence for a direct, content-specific involvement of the “front” of the cortex, including most prefrontal regions, is missing or unclear” (Boly *et al.*, 2017), while others argued that “the literature highlights prefrontal cortex’s essential role in enabling the subjective experience in perception” (Odegaard, Knight and Lau, 2017). Although this issue remains unresolved, the debate reflects how different theories of consciousness entail different hypotheses about which brain regions contribute directly to experience. Hence, evidence showing regional differences in contribution to the capacity for, or contents of, consciousness may provide specific empirical support for some theories over others.

While frontal parts of the cortex have been suggested to be crucial for consciousness (Del Cul *et al.*, 2009), others have pointed to evidence that apparent frontal involvement in consciousness is likely confounded by task-related processes such as working memory, attention, or preparation for motor response (Boly *et al.*, 2017). In general, behavioural responsiveness has often been confounded with consciousness (Sanders *et al.*, 2012). Ability to respond coherently to external stimuli is used as the main criterion for determining whether non-communicating patients and non-human animals are conscious (Chernik *et al.*, 1990; Giacino, Kalmar and Whyte, 2004; Gao and Calderon, 2020). However, this approach is based on the assumption that unresponsive states are always unconscious, which is at odds with evidence that vivid experiences can occur in unresponsive states. For instance, dreams can occur in all stages of sleep (Nielsen, 2000) and during general anaesthesia (Noreika *et al.*, 2011). Furthermore, patients can, be conscious but unresponsive for decades after brain damage (Vanhaudenhuyse *et al.*, 2018), or they can be painfully aware during general anaesthesia for surgery, while assumed to be unconscious (Ghoneim *et al.*, 2009). Conversely, quite complex behaviours can be preserved during conditions that are typically assumed to be unconscious, e.g. sleepwalking, sleep talking, and presumed unconscious behaviours during certain epileptic seizures, and in unresponsive wakefulness syndrome (Blumenfeld, 2005; Laureys *et al.*, 2010). Thus, it is essential to distinguish brain regions necessary for consciousness from those that are necessary for behavioural responsiveness.

Recently, several measures aiming to objectively assess global states of consciousness independently of motor or sensory functions have been developed. Specifically, the perturbational complexity index (PCI; Casali *et al.*, 2013), and the more general measure PCI^ST^ (“PCI-state transition”; Comolatti *et al.*, 2019), have been shown to reliably and consistently assess the state of consciousness in humans in accordance with the subjects’ immediate or delayed reports of experience (Sarasso et al. 2015; Casarotto et al. 2016; Rosanova et al. 2018). Recently, PCI^ST^ has also been shown to work consistently in rodents undergoing propofol, sevoflurane, and ketamine anaesthesia (Arena *et al.*, 2020). PCI and PCI^ST^ quantify the spatiotemporal complexity of repeatable, global, cortical, electrophysiological responses to a local, direct cortical stimulation, thus estimating how much the resulting deterministic neuronal activations are both integrated and differentiated across cortical areas and time. This measure was inspired by the idea that highly complex interactions are fundamentally associated to the conscious experience, as proposed by the Integrated Information Theory of consciousness (IIT; Tononi, 2004; Massimini et al., 2009). The PCI measure is also coherent with predictions of the Global Neuronal Workspace Theory (GNW) of consciousness, which suggests that conscious processing is associated with global and durable “broadcasting” of information (Mashour et al., 2020). Hence, PCI may be considered a candidate index of consciousness even in cases where we do not know the ground truth (Casarotto et al. 2016; Comanducci et al. 2020). Therefore, for the remainder of this paper, we interpret a significant drop in PCI from what is observed in wakefulness as indicative of a relative reduction in capacity for consciousness.

While PCI is an index of the capacity for consciousness based on evoked cortical dynamics, measures that quantify spectral properties of spontaneous cortical activity have also successfully been used for a long time to study brain states (Loomis, Harvey and Hobart, 1937; Fernandez *et al.*, 2017; Siclari *et al.*, 2017; Colombo *et al.*, 2019; Lendner *et al.*, 2020). Specifically, during NREM sleep and general anaesthesia, both electroencephalography (EEG) and local field potentials (LFP) are characterized by high amplitude, low frequency (LF, ≤ 4 Hz) oscillations or slow-waves (Massimini *et al.*, 2004; Vyazovskiy *et al.*, 2009; Brown, Lydic and Schiff, 2010). This slow-wave activity reflects a bistable network dynamic, where neurons synchronously alternate between an up-state, with depolarized membrane potential and firing, and a down-state with neuronal hyperpolarization and silence (Steriade, Nuñez and Amzica, 1993; Steriade, Timofeev and Grenier, 2001; Volgushev *et al.*, 2006; Vyazovskiy *et al.*, 2009), possibly due to increased inhibition, adaptation, and synaptic fatigue (Steriade, Timofeev and Grenier, 2001; Compte *et al.*, 2003; Esser, Hill and Tononi, 2007; Funk *et al.*, 2017). Conversely, during wakefulness the EEG is mainly characterized by low amplitude, high frequency (HF, ≥ 20 Hz) oscillations, which reflect tonic neuronal depolarization and firing (Steriade, Amzica and Contreras, 1996; Steriade, Timofeev and Grenier, 2001; Mukovski *et al.*, 2007; Vyazovskiy *et al.*, 2009). This has inspired several measures for quantifying the cortical state of activation based on the relation between high and low frequency EEG power (Mukovski *et al.*, 2007; Fernandez *et al.*, 2017; Gao, Peterson and Voytek, 2017; Siclari *et al.*, 2017; Colombo *et al.*, 2019; Lendner *et al.*, 2020). Thus, the HF/LF (power) ratio tends to drop when there is more inhibition and deactivation, whereas a higher HF/LF ratio suggests neuronal activation (Mukovski et al. 2007; Poulet and Crochet 2018; Gao et al. 2017; Lombardi et al. 2017). Coherently, the spectral relation between HF and LF powers has been associated with the excitation/inhibition balance (Gao, Peterson and Voytek, 2017) and the ratio between low and high frequency powers was found to correlate with motor activity in mice (Fernandez *et al.*, 2017). Thus, local changes in HF/LF ratio may indicate changes in cortical activation (Poulet and Crochet, 2018).

While measures of consciousness typically evaluate global brain dynamics, specific and localized changes in the HF/LF ratio might occur within the same brain state, and might affect behaviour or conscious experience. For example, local cortical sleep, which can affect task performance in rats, involves localized slow waves and neuronal silence in an otherwise awake brain state, dominated by low amplitude and fast oscillations, with underlying tonic neuronal firing (Vyazovskiy *et al.*, 2011). Furthermore, posterior increases in HF/LF ratio during NREM sleep successfully predicted whether or not humans reported dreaming (Siclari *et al.*, 2017). Thus, HF/LF can vary regionally within global states, but it is not known whether local reductions in HF/LF are associated with reduced consciousness level. More generally, it is still unknown whether any particular localized spectral properties are related to the brain’s global capacity to sustain consciousness.

In this study, we aimed at investigating this relation, asking whether specific regional changes of cortical activation, assessed by the HF/LF ratio, are associated with changes in capacity for consciousness, assessed by PCI^ST^. By using two distinct levels of ketamine anaesthesia, we dissociated levels of behavioural responsiveness (assessed by responses to pain stimuli) from the capacity for consciousness, and investigate whether regional HF/LF ratio reliably covaries with changes in PCI^ST^. The results may be used to validate predictions from theories of consciousness by constraining which cortical regions mainly underlie consciousness as opposed to behavioural responsiveness.

## MATERIALS AND METHODS

### Animal model and experimental data

Six adult, male, Sprague-Dawley rats (n = 6; body weight ~370 g) were used in this study. All the experiments and animal care procedures were conducted at the University of Oslo and were approved by the Norwegian Authority, Mattilsynet (FOTS: 11812) in agreement with Norwegian law of animal handling. Efforts were made to avoid or minimize animals’ pain and distress. Rats were caged in enriched environments, with *ad libitum* access to food and water and were exposed to a 12:12 hour light–dark cycle at 23°C constant room temperature.

In addition to new data from six rats, we also used data from a set of previously published experiments (Arena, Thon and Storm, 2019), in which we recorded epidural EEG continuously and in response to electrical stimulation of secondary motor cortex in 12 head-restricted rats (Arena *et al.*, 2020). For 8 of the rats, the recordings were performed during wakefulness and subsequent light ketamine anaesthesia. In 6 of these 8 rats, a third round of recording was also performed during an increased dosage of ketamine anaesthesia, given at the end of the 6 experiments. Here, we report results from this subset of 6 animals, based on new analysis of data from wakefulness and light ketamine anaesthesia and compared with new data from deep ketamine anaesthesia. All the 6 rats were exposed to all conditions, allowing internal controls.

### Experimental procedure

Epidural EEG was recorded by a grid of 16 screw electrodes (stainless steel, 1.2 mm calibre), which were chronically implanted through the skull, in contact with the dura. The recording electrodes were organized symmetrically with respect to the sagittal suture and spanned most of the cortical surface of both hemispheres (**Supplementary Fig.1**). Bilateral frontal cortex was covered by 6 electrodes, named M2_R_, M2_C_ and M2/M1 that medially overlaid the rostral and caudal part of the secondary motor cortex and part of the primary motor cortex, respectively. Other 6 electrodes covered left and right parieto-occipital associative cortices: PA electrodes covered lateral parietal cortex, RS/PA medially covered retrosplenial and parietal cortex, while RS/V2 electrodes covered the posteromedial cortex, over the caudal part of retrosplenial cortex and medial part of secondary visual cortex. The last 4 electrodes, bilateral S1 and V1, overlaid the primary somatosensory cortex and the primary visual cortex respectively. Event-related potentials (ERPs) were recorded in response to electrical stimulation of the right secondary motor cortex by a bipolar tungsten electrode (see **Supplementary Fig. 1** for detailed electrode locations with respect to bregma; Paxinos & Watson, 2007). Standard surgical procedure under a regime of controlled general anaesthesia/analgesia was adopted for implantation of chronic electrodes, and after 3 days of recovery, rats were habituated to head and body restriction in at least 3 subsequent days, as previously described (Arena *et al.*, 2020). The electrophysiological recording/stimulation began only when rats did not show any sign of distress and were calm within the recording setup, with the head connected to a fixed head-bar by two chronically implanted clamps and with the body inserted in a transparent acrylic tube, with a natural posture. The tail was left outside the tube to test reflex motor responses to pain stimulation.

The 6 rats were subjected to electrophysiological recording/stimulation sessions during wakefulness, and subsequent ketamine anaesthesia. Ketamine (Vetoquinol, Ittigen, Switzerland) was infused at a constant rate via a 26 GA catheter in the tail vein. Subcutaneous injection of glycopyrrolate 0.01 mg/kg was also performed to reduce the increased salivation, eye ointment was applied to maintain eyes humid and body temperature was kept at 36.5 - 37.5 °C by a heating pad, as previously described in detail. During the recording session the stimulating electrode was connected to an isolated current stimulator (Isolator HG203, High Medical, London Uk) triggered by a voltage pulse generator (2100, A-M System, Washington DC, USA), while the epidural EEG electrodes were connected to a 16-channel unipolar amplifier referenced to ground (RHD2132, Intan Technologies, Los Angeles, CA, USA), and controlled by Open Ephys system (Siegle *et al.*, 2017), which acquired and digitized the electrophysiological signal at 10 or 30 kHz, 16-bit resolution.

The EEG activity was continuously recorded from all 16 channels, in a dark environment, in which all rats received ~100 electrical monophasic current pulses of 50 μA, 1 ms, delivered at 0.1 Hz, at first during wakefulness (W), and during subsequent ketamine anaesthesia. The anaesthesia was induced by an intravenous (iv) bolus injection of ketamine 30 mg/kg, and then maintained at the initial constant rate of ketamine 1.75 mg/kg/min iv, by a syringe pump. After 10 min from induction, the tail was pinched 3 times by a forceps to check the presence of behavioural reaction to pain stimulation. The behavioural response typically consisted in a lateral and wide movement of the tail and/or in the alteration of the respiratory rhythm, with the occurrence of a deeper breath, accompanied by a sudden chest movement. The infusion rate was stepwise increased by adding 4 % of the initial dosage until the behavioural response was absent (3 min between each increment). The resulting minimal dosage of ketamine that abolished behavioural reaction to pain stimulation was 1.78 ± 0.02 mg/kg/min, iv (mean ± SEM across rats; 1.8 mg/kg/min for simplicity) and defined the condition “ketamine 1” or K1. The same recording/stimulation was also repeated afterwards, during deeper ketamine anaesthesia (“ketamine 2” or K2), at the constant rate of 3.5 mg/kg/min iv, which corresponded to 2 times the initial constant rate.

### Analysis of electrophysiological signal

The acquired electrophysiological data were analysed in MATLAB2016a (Math Works, Natick, Massachusetts, USA) and Origin 9.1 (OriginLab, Northampton, Massachusetts, USA) and pre-processed as described before (Arena *et al.*, 2020). Raw epidural EEG was band pass filtered (0.5-80 Hz, butterworth, 3rd order), down-sampled to 500 Hz and ERP epochs of 10 s were extracted for each channel, centred at the stimulus onset (from −5 to 5 s). All epochs were offset corrected and trials with high voltage artefacts in their baseline were removed. The first 90 trials of pre-processed signal were used for analysis.

The cortical excitation in response to electrical stimulation was quantified by the root mean squared (rms) amplitude of the first 50 ms of the mean ERP for all electrodes, and then averaged across electrodes. Spectral powers and phases of ERPs were obtained from Morlet wavelet convolution (3 cycles wavelets, linearly spanning from 1 to 80 Hz, with 1 Hz resolution), which was performed for each trial and channel. The spectral powers were normalized over the mean power of the baseline (−500 to −200 ms) across trials, for each respective frequency and channel. The mean relative powers across trials were then dB converted and bootstrap statistic (500 permutations; positive and negative thresholds based on the obtained distribution of the maximum and minimum dB values in the baseline window, α = 0.05) was applied for each frequency and channel to conserve only the significant dB variations from respective baseline. The resulting relative spectral power was averaged across trials in the high frequency range (HF, 20-80 Hz) to estimate the temporal dynamic of the neuronal activation underlying the EEG signal (Mukovski *et al.*, 2007; Pigorini *et al.*, 2015; Rosanova *et al.*, 2018; Arena *et al.*, 2020). Late increments of HF power (> 0 dB) were detected in a time window from 80 to 800 ms (Arena *et al.*, 2020). The inter-trial phase clustering (ITPC; Cohen, 2014) was computed for each frequency and channel and bootstrap statistic (500 permutations; threshold based on the obtained distribution of the maximum ITPC values in the baseline window, α = 0.01) was used to conserve only the ITPC increments that differed significantly from the respective baseline (from −500 to −200 ms). The time point of the last significant ITPC value within 800 ms after stimulation and in a broad frequency range (5-80 Hz) was detected for each channel and considered to be the duration of the phase-locked EEG response, which quantified the temporal extension of the deterministic effect of the electrical stimulation (ITPC drop time; David, Kilner and Friston, 2006; Pigorini *et al.*, 2015; Rosanova *et al.*, 2018; Arena *et al.*, 2020). PCI^ST^ was used to estimate the capacity for consciousness (Comolatti *et al.*, 2019; Arena *et al.*, 2020) and was assessed in the full ERP window, from 0 to 600 ms, and across time, in short moving windows of 100 ms, with 50 ms of overlap. PCI^ST^ was computed using the available code at [github.com/renzocom/PCIst] with the same parameters previously described (Comolatti *et al.*, 2019; Arena *et al.*, 2020). The functional connectivity across cortical areas in response to stimulation was also used to estimate the level of integration and differentiation of the cortical network, as previously described in detail (Arena *et al.*, 2020). The inter-site phase clustering (ISPC) was assessed as the consistency across trials of the phase difference across channels for all time-frequency points (Cohen, 2014). ISPC was calculated for each channel pair, the respective mean ISPC of the baseline (from −500 to −200 ms) was subtracted and bootstrap statistic (500 permutations; positive and negative thresholds based on the obtained distribution of the maximum and minimum ISPC values in the baseline window, α = 0.05) was adopted to conserve only the significant variations from baseline. The ISPC values that could be determined by volume conduction (clustering around 0 or pi) were set to 0 and the remaining ISPC values were averaged in the frequency range 5-14 Hz and between 180 and 400 ms. The proportion of the number of mean positive ISPC values for each channel was defined as the connectivity degree (CD) of the electrode (Cohen, 2014; Arena *et al.*, 2020).

The spontaneous cortical activity associated with the different experimental conditions was quantified from 90 epochs of 5 seconds of epidural EEG signal that preceded the electrical stimulation (from −5 to 0 s). A Morlet wavelet convolution (6 cycles, 80 wavelets, linearly spanning from 1 to 80 Hz) was performed on each epoch for all channels. Spectral powers were extracted and averaged across samples, trials and channels, obtaining a global estimation of the power of each frequency for each animal and condition. The resulting periodogram was linearly fitted in Log-Log coordinates, in the frequency range 20-40 Hz. The slope of the obtained linear function was the spectral exponent of the 1/f function and was used to quantify the distribution of frequency powers in the spontaneous EEG activity (Gao, Peterson and Voytek, 2017; Colombo *et al.*, 2019; Arena *et al.*, 2020; Lendner *et al.*, 2020). Instantaneous powers were normalized by 1 and converted in dB, and also averaged in high and low frequency ranges (HF 20-80 Hz, LF 1-4 Hz respectively), across and for each single channel, before dB conversion. The ratio between HF power and LF power (HF/LF ratio) was also computed to estimate the level of activation for the cortical area underlying each channel (Fernandez *et al.*, 2017; Poulet and Crochet, 2018) and its contribution in cortical complexity and capacity for consciousness (Siclari *et al., 2017*).

### Statistics

All results are expressed as mean ± SEM and error bars and shades represent SEM in figures. Shades in linear regressions represent the 95% confidence band. The topographical plots in the figures report the Laplace interpolation of a variable over the dorsal surface of a skull, anchored to the true electrode locations. The function of the colour maps is only for a better visualization, since all the analysis were performed at the level of single channels. Parametric statistics were adopted after assessing the normality of distribution of the measured variables, by applying the Shapiro-Wilk test, in a population of 12-14 rats during wakefulness. All the variables were tested in a repeated measure design. Thus, principal and interaction effects were tested with one-way or two-ways repeated measures ANOVA (rANOVA), in which Greenhouse–Geisser correction was applied when sphericity could not be assumed, while group comparisons were tested with Student’s paired-samples t-test. Because one channel was removed from the analysis in 2 rats, two-way ANOVA was used to compare the spatial distribution of variables between left and right hemispheres. Linear fitting was performed with the least-square method. To evaluate correlations and goodness of fit, the coefficient of determination, R^2^, was computed and t-test was performed to test the null hypothesis of slope = 0, establishing a *P* value. When multiple hypotheses were tested across conditions or along cortical areas, the Bonferroni–Holm correction was adopted (number of conditions = 3, corrected α = 0.01666; number of cortical areas = 8, corrected α = 0.00625 for each hemisphere). Gaussian v-test was used to test volume conduction in connectivity analysis (Cohen, 2014; Arena *et al.*, 2020). All statistics are two-tailed. The statistical significance in figures are represented as follows: *P* < 0.05 *, *P* < 0.01 **, *P* < 0.001 ***, *P* ≥ 0.05 ns (not significant).

## RESULTS

### Perturbational complexity was independent of behavioural responsiveness, but was reduced by increasing ketamine dosage

In previous studies in humans, ketamine has been found to induce unresponsiveness combined with vivid, dream-like experience and high cortical complexity (Sarasso et al., 2015). Thus, to dissociate cortical complexity from behavioural responsiveness, we recorded spontaneous and evoked EEG activity in 6 rats, during wakefulness (W) and subsequent constant intravenous infusion of ketamine. We also repeated the electrophysiological experiment with an increased administration rate, to test for a possible dosage dependency. As reported previously (Arena *et al.*, 2020), we adjusted the first, low ketamine infusion rate (K1) for each single rat, to the minimal dose that induced behavioural unresponsiveness (i.e. no motor response to pain stimulation). This first dosage, K1, was approximately 1.8 mg/kg/min, iv, while the second subsequent ketamine dose, K2, was set to about two times the first one, at 3.5 mg/kg/min, iv.

During wakefulness, the spontaneous EEG activity was characterized by low amplitude, fast oscillations, typical of cortical activation (**Fig. 1A**). Interestingly, despite loss of behavioural responsiveness, the fast oscillations persisted in both K1 and K2 conditions (**Fig. 1A**), and a general increase of EEG amplitude occurred with ketamine infusion, as shown by the averaged periodogram across channels, which scaled up from wakefulness to K1 and further to the K2 condition (**Fig. 1B**). Likewise, the mean HF power (20-80 Hz) across channels increased from wakefulness to K1 and further to K2 (**Fig. 1C**; W: 42.42 ± 0.54 dB, K1: 47.52 ± 0.83 dB, K2: 49.15 ± 0.64 dB; one-way rANOVA, principal effect of condition, *P* = 9.1942*10-8; paired samples t-test, W vs K1, *P* = 0.0005, W vs K2, *P* = 6.8217*10^−5^, K1 vs K2, *P* = 0.0032). As for HF activity, also the mean LF (1-4 Hz) power increased from wakefulness to K1 and further to K2 (**Fig. 1D**; W: 73.57 ± 1.26 dB, K1: 77.75 ± 0.72 dB, K2: 79.75 ± 0.54 dB; one-way rANOVA, principal effect of condition, *P* = 0.0021; paired samples t-test, W vs K1, *P* = 0.0440, W vs K2, *P* = 0.0271, K1 vs K2, *P* = 0.0271). The spectral exponent of the mean periodograms was similar across conditions, indicating a general scaling across all frequencies (**Fig. 1E**; W: −1.82 ± 0.17, K1: −1.41 ± 0.12, K2: −1.57 ± 0.12; one-way rANOVA, principal effect of condition, *P* = 0.1959; paired samples t-test, W vs K1, *P* = 0.1719, W vs K2, *P* = 0.3077, K1 vs K2, *P* = 0.2838).

**Fig. 1.**
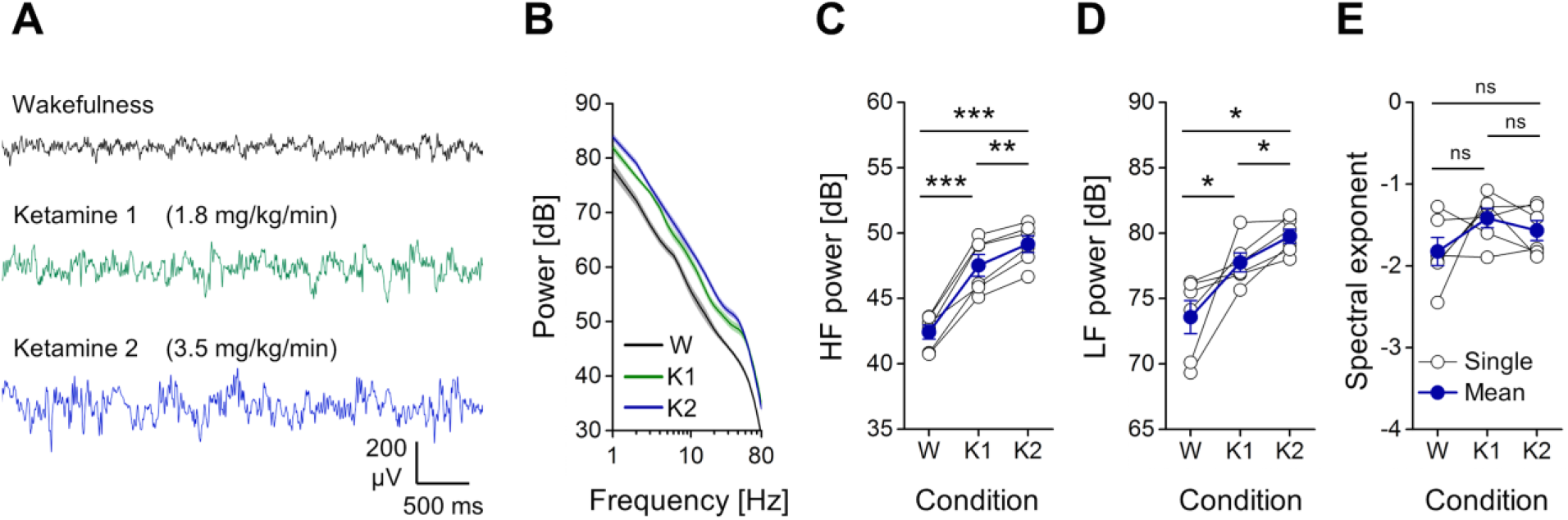
The averaged power spectrum of spontaneous EEG was scaled up from wakefulness to ketamine anaesthesia and by increasing ketamine dosage. **(A)** Example of spontaneous EEG (5 seconds) from the retrosplenial/parietal cortex (RS/PA) of one rat during wakefulness (W), light ketamine anaesthesia (ketamine 1, K1; administration rate: 1.8 mg/kg/min i.v.) and deep ketamine anaesthesia (ketamine 2, K2; administration rate: 3.5 mg/kg/min i.v.). **(B)** Mean periodograms from 16 channels and 6 rats exposed to the same conditions of A (shades represent SEM across rats). **(C)** Variations of mean high frequency power (HF, 20-80 Hz) and **(D)** low frequency power (LF, 1-4 Hz) are shown for each rat across conditions and increased from W to K1 and from K1 to K2. **(E)** The spectral exponents of the averaged periodograms across channels are also reported for each rat and condition.

Across conditions, the same electrical stimulation induced a similar initial cortical excitation, quantified by the rms amplitude of the early ERP deflections (**Fig. 2A, B**; W: 52.63 ± 5.13 μV, K1: 47.65 ± 3.91 μV, K2: 44.84 ± 4.31 μV; one-way rANOVA, principal effect of condition, *P* = 0.1943; paired samples t-test, W vs K1: *P* = 0.2942, W vs K2: *P* = 0.1613, K1 vs K2: *P* = 0.1132). Consistently, the electrical pulse evoked an early, broad-band increment in power that was similar across conditions. This activation was quickly followed by a period of relaxation, with HF activity similar to or below the baseline (dB ≤ 0), which lasted 150.52 ± 12.04 ms, averaged across rats and conditions (**Fig. 2A**). However, after this first dynamic, the ERP developed differently through time in the different conditions. During wakefulness and with low ketamine dose, the ERPs showed long-lasting waveforms that were phase-locked across trials, thus still deterministically caused by the stimulation. Conversely, with the high ketamine dosage, the phase-locked response quickly died out, indicated by an earlier ITPC drop, averaged across channels (**Fig. 2A, C**; W: 347.35 ± 18.85 ms, K1: 369.76 ± 59.86 ms, K2: 152.32 ± 24.17 ms; one-way rANOVA, principal effect of condition, *P* = 0.001; paired samples t-test, W vs K1: *P* = 0.7068, W vs K2: *P* = 0.0014, K1 vs K2: *P* = 0.01). The long-lasting response in wakefulness and K1 also showed a later increase of HF power in most of the channels (W: 100 % of channels; K1: 92.36 ± 4.33 % of channels). Such HF activation was largely associated with the deterministic response, since its onset preceded the ITPC drop, as shown by comparing the times of these events from the electrodes with such late increase in HF power (**Fig. 2A, D**; paired samples t-test, ITPC drop time vs Late HF power onset; in W: n = 94 channels, *P* = 7.8199*10^−45^, in K1: n = 87 channels, *P* = 2.6807*10^−13^). In contrast, with high ketamine dose, a late HF power activation was still detected in some electrodes (46.6 ± 14.93 % of channels), but was not phase-locked, as it occurred after the ITPC drop (**Fig. 2A, D**; paired samples t-test, ITPC drop time vs Late HF power onset, in K2: n = 44 channels, *P* = 2.1199*10^−5^). These results were also associated with an overall reduction of the functional connectivity across channels in the K2 condition, which also corresponded to a reduced diversity of cortical connectivity (**Supplementary Fig. 2**). Coherently, K2 changed the time course of PCI^ST^, which was initially high and quickly decayed soon after the stimulation in all conditions. However in wakefulness and with low ketamine dose, PCI^ST^ built up again, reaching similar values after 200 ms. Then PCI^ST^ dropped again, faster in K1 than W, until fading. Conversely, with high ketamine dose, PCI^ST^ never recovered after the early decay and remained close to 0 (**Fig. 2A, E**). The time course was in line with the perturbational complexity calculated for the entire response window (0-600 ms), thus PCI^ST^ showed slightly higher values in wakefulness than in K1 condition, but the difference was not statistically significant for this sample (although a significant difference was found for a larger data set; Arena *et al.*, 2020, see Discussion). With high ketamine dose, however, PCI^ST^ was significantly lower than the other conditions (**Fig. 2E;** W: 78.47 ± 8.44, K1: 59.57 ± 5.77, K2: 23.75 ± 6.55; one-way rANOVA, principal effect of condition, *P* = 4.3026*10^−5^; paired samples t-test, W vs K1, *P* = 0.0535, W vs K2, *P* = 0.0028, K1 vs K2, *P* = 0.0005).

**Fig. 2.**
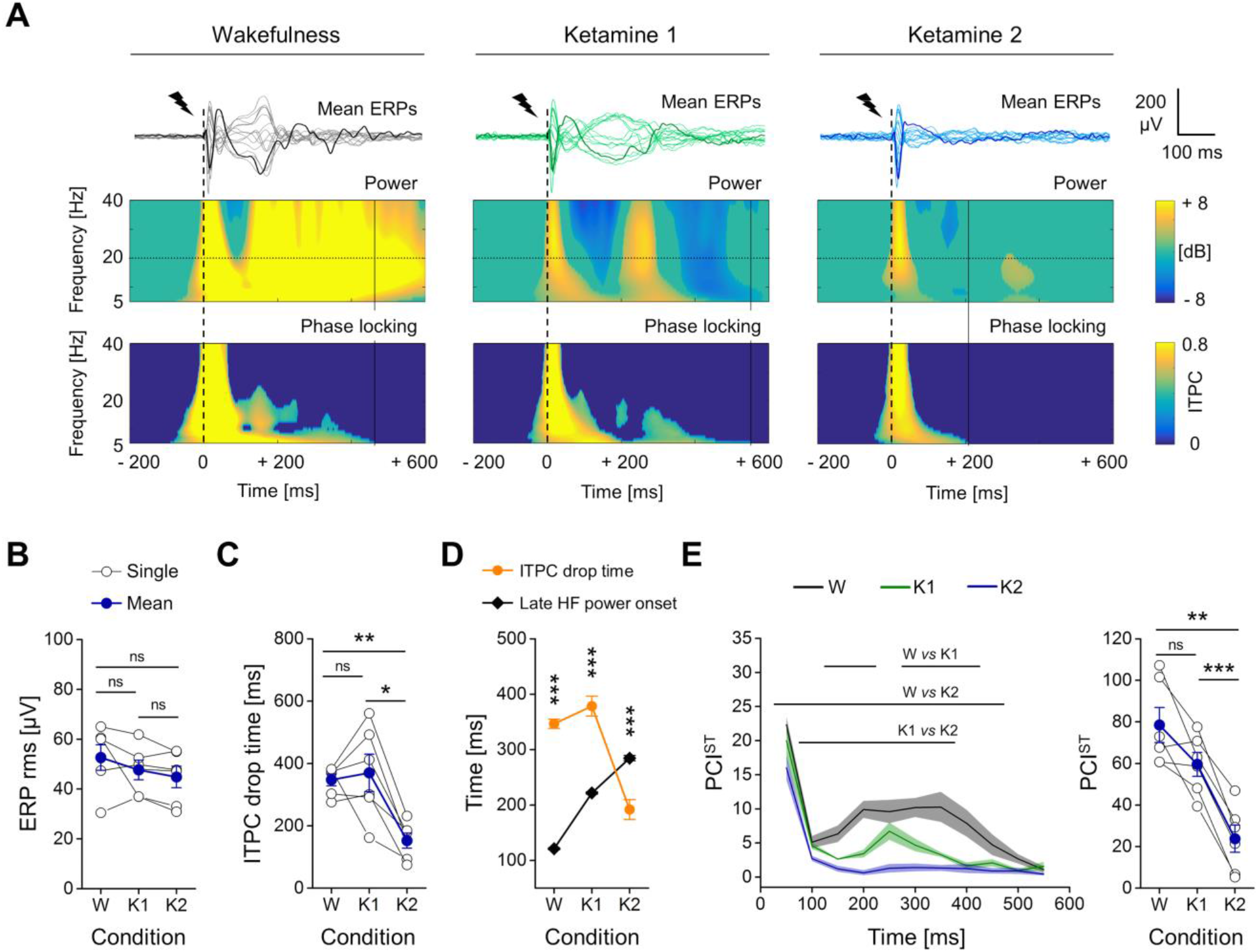
The spatiotemporal dynamics of ERPs revealed a drop in complexity from low to high dosage of ketamine. **(A)** *Top,* superimposition of mean ERPs from all 16 electrodes in response to single pulse stimulation (1 ms, 50 μA; dashed line) of the right secondary motor cortex (M2), from the same rat during wakefulness (W, *left*) and light and deep ketamine anaesthesia (K1, *middle* and K2, *right* respectively). One averaged ERP from the same channel over the right primary somatosensory cortex (S1) is in bold for clarity. *Middle,* power spectrogram (dB) and, *bottom,* phase-locking across trials (ITPC) from the channel shown in bold above (right S1). The ITPC drop time is indicated by vertical continuous lines. **(B)** The rms amplitude of the early ERP (first 50 ms from stimulus onset) averaged across channels is shown for all rats and conditions. **(C)** The ITPC drop time (in frequency range 5-80 Hz) averaged across channels is plotted for each rat and condition. **(D)** The ITPC drop time and the onset of later increased HF power were averaged across channels from all rats and shown for each condition. **(E)** *Left*, time courses of mean PCI^ST^ (moving windows of 100 ms, 50 ms overlap) and standard errors (shaded) are plotted for all conditions (horizontal lines indicate periods of statistically significant difference between conditions, *P* < 0.05). *Right*, PCI^ST^ in range 0-600 ms is shown for each rat and condition.

### The strong reduction in perturbational complexity was associated with a selective deactivation of bilateral posteromedial cortex

We found that high-dose ketamine caused a non-linear, dose-dependent drop of PCI^ST^ (**Fig. 2**), but this drop was not associated with a clear overall transition of spontaneous EEG towards slow waves and reduced HF power (**Fig. 1**). Thus, we investigated the spectral features of spontaneous activity from single cortical areas with higher detail, by computing the local, instantaneous HF/LF ratio (Fernandez *et al.*, 2017; Siclari *et al.*, 2017; Poulet and Crochet, 2018; see **Fig. 3A**). By averaging across time (from −5 to 0 s) and trials for each electrode, we obtained a topographical distribution of the HF/LF ratio, which was similar between left and right hemispheres, and this symmetry persisted in all conditions (**Fig. 3B;** two-way ANOVA, principal effect of lateralization, in W: *P* = 0.6934, in K1: *P* = 0.9902, in K2: *P* = 0.9789). By comparing the mean HF/LF ratio across conditions, for each cortical area, we found that it significantly decreased from wakefulness to low ketamine dosage only in right M2/M1 and S1 cortex, although weak, non-significant reductions were also seen elsewhere (**Fig. 3B**). In contrast, high dose of ketamine selectively reduced the mean HF/LF ratio in the bilateral posteromedial cortex (left and right RS/V2 channels), indicating a specific deactivation of this part of the cortex, with respect to the K1 condition (**Fig. 3A, B**, **Supplementary Fig. 3**). In line with these results, the HF/LF ratios at both bilateral RS/V2 and right M2/M1 cortex were also reduced from wakefulness to the K2 condition (**Fig. 3B**). To test whether the level of activation of any cortical area was effectively able to predict the complexity of global cortical dynamics, we assessed possible regional correlations between HF/LF ratio and PCI^ST^ (**Fig. 3C**). We found that the HF/LF ratio from spontaneous activity of bilateral posteromedial cortex (left, right RS/V2) was highly, linearly correlated with the PCI^ST^ value across conditions (**Fig. 3C, D;** left RS/V2: linear fit, R^2^ = 0.590, *P* = 0.0016; right RS/V2: linear fit, R^2^ = 0.568, *P* = 0.0024). A weaker but significant correlation was also identified only at the level of the right secondary motor cortex (**Fig. 3C, D;** right M2_C_: linear fit, R^2^ = 0.417, *P* = 0.0264).

**Fig. 3.**
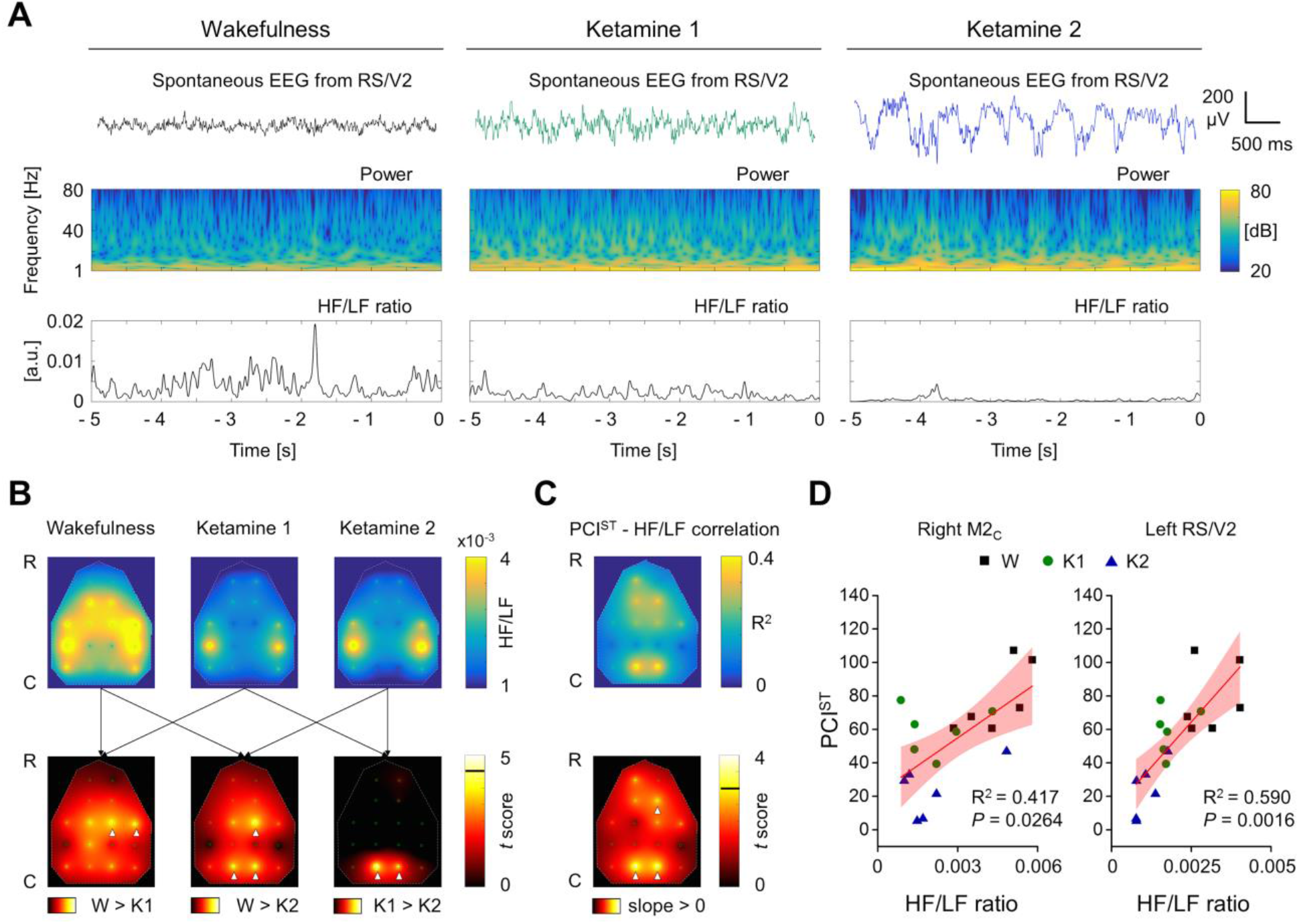
The HF/LF ratio of spontaneous EEG in bilateral posteromedial cortex was selectively reduced by increasing ketamine dosage and correlated with the level of PCI^ST^. **(A)** *Up*, Example of spontaneous EEG (5 seconds) from the posteromedial cortex (RS/V2) of one rat during wakefulness (W), light ketamine anaesthesia (ketamine 1, K1) and deep ketamine anaesthesia (ketamine 2, K2) and *below*, the relative spectrogram and the ratio between high frequency (HF, 20-80 Hz) and low frequency (LF, 1-4 Hz) powers (HF/LF ratio) in time. **(B)** The colour maps show the topographical distributions (R-C: rostral-caudal) of the 16 EEG electrodes (small green circles) and, *above*, the spatial interpolations of the HF/LF ratio, averaged across time, trials and rats, for each condition. *Below*, the colour maps report the spatial interpolation of the *t* scores (paired samples t-test) from comparing the HF/LF ratio across conditions for each channel (*left*, wakefulness vs ketamine 1; *middle*, wakefulness vs ketamine 2; *right*, ketamine 1 vs ketamine 2). The horizontal black line in the colour bar indicates the threshold for statistical significance (*t*_*5*_ = 4.5258, Bonferroni-Holm corrected). White arrowheads indicate the channels with statistically significant difference across conditions. **(C)** *Above,* the colour map shows the spatial distribution of the coefficient of determination R^2^ from the correlation between PCI^ST^ and HF/LF ratio, across rats and conditions, for each channel. *Below*, the colour map reports the spatial interpolation of *t* scores, assessing the statistical significance of the correlation for each channel. The horizontal black line in the colour bar indicates the threshold for statistical significance (*t*_*16*_ = 3.1458, Bonferroni-Holm corrected). White arrowheads indicate the channels showing statistically significant correlations. The correlations of right M2_C_ and left RS/V2 are reported in **(D)** with respective R^2^ and *P* values.

Next, we assessed the same correlations between local HF/LF ratio and global PCI^ST^, this time distinguishing between brain states and state transitions. We first evaluated the correlation in wakefulness condition alone, where both behavioural responsiveness and high perturbational complexity were present (**Fig 4A, B;** data from 3 different recordings, performed in 3 different days on the same rats were considered only in this condition, to increase the number of observations). Then, we repeated the estimation of the correlations by considering the conditions of wakefulness and low dosage of ketamine, when perturbational complexity is still high, but behaviour transitions from responsiveness to unresponsiveness (**Fig. 4C, D**). Finally, we evaluated the correlations in conditions of low and high ketamine dosages, when only the variation of perturbational complexity occurred, within the same unresponsive behavioural state (**Fig. 4E, F**). With this, we attempted to identify possible roles that specific cortical regions might have in specific state transitions. Within the wakefulness condition, we could not identify any significant correlation between PCI^ST^ and HF/LF ratio at the level of any cortical area. Nevertheless, the correlations with higher R^2^, which were also closer to the threshold for statistical significance, seemed to be clustered in secondary motor cortex (**Fig 4A, B**; right M2_C_: linear fit, R^2^ = 0.377, *P* = 0.0538; left RS/V2: linear fit, R^2^ = 0.077, *P* = 1). Likewise, the HF/LF ratio could not clearly predict PCI^ST^ between wakefulness and low ketamine dose, when only behavioural responsiveness changed, for any of the cortical areas (**Fig 4C, D;** right M2_C_: linear fit, R^2^ = 0.386, *P* = 0.2173; left RS/V2: linear fit, R^2^ = 0.271, *P* = 0.4124). On the other hand, by considering only the variations induced by increasing ketamine dosage (K1 and K2), we found a significant and strong correlation selectively associated to the left RS/V2 cortex, thus indicating that the state of activation or deactivation of posteromedial cortex could effectively linearly predict the complexity level of the entire cortical network and its breakdown (**Fig 4E, F;** right M2_C_: linear fit, R^2^ = 0.064, *P* = 0.1; left RS/V2: linear fit, R^2^ = 0.568, *P* = 0.0370). Coherently with these results, by considering only the conditions of wakefulness and high dose of ketamine, the variation of behavioural responsiveness could not be separated from changes in perturbational complexity, hence a significant correlation between PCI^ST^ and the HF/LF ratio was detected in both frontal cortex and posteromedial cortex (**Supplementary Fig. 4**).

**Fig. 4.**
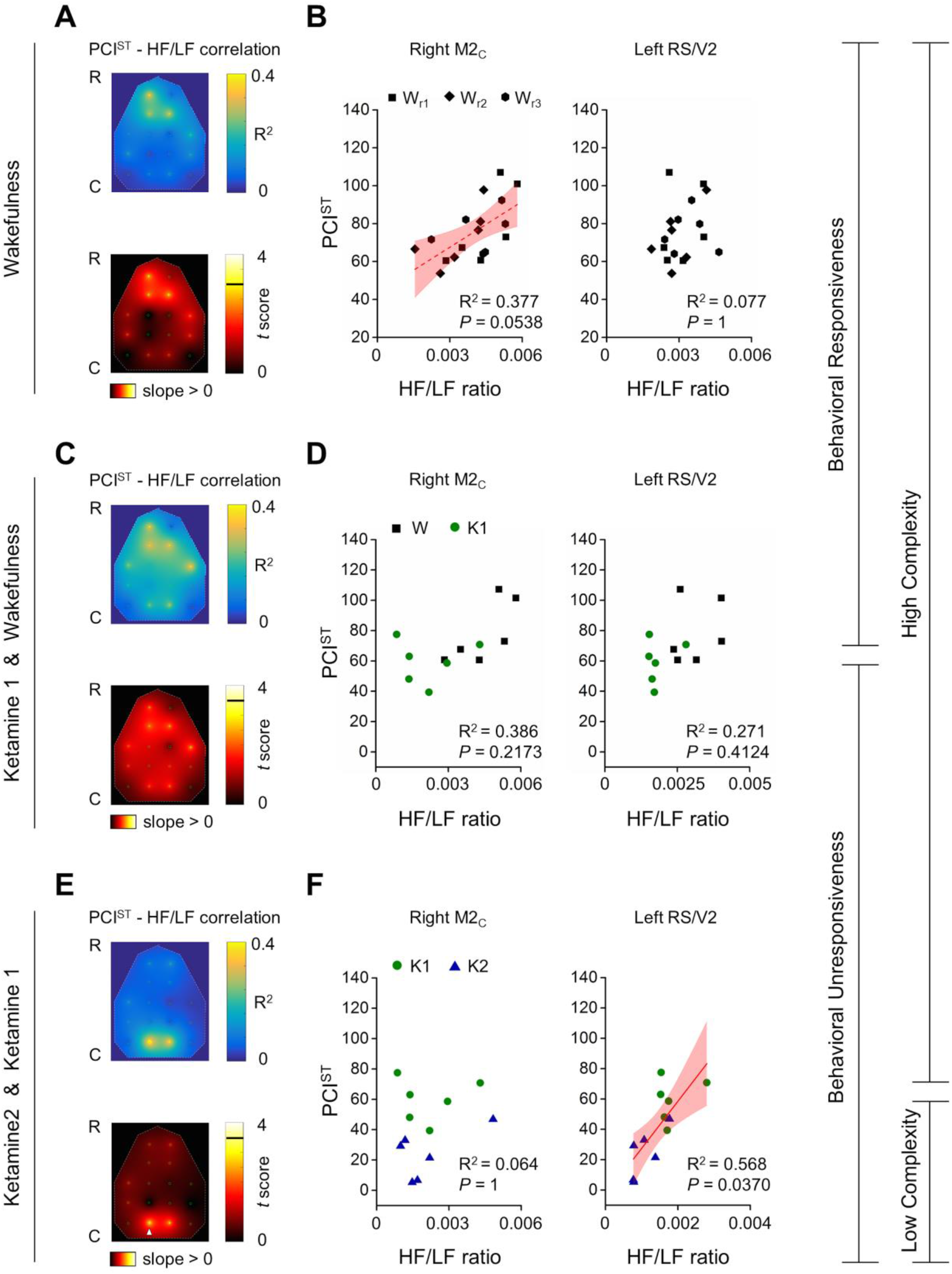
HF/LF ratio of spontaneous EEG from posteromedial cortex selectively correlated with PCI^ST^ in conditions of behavioural unresponsiveness, with light and high ketamine anaesthesia. **(A, B)** The putative correlations between PCI^ST^ and HF/LF ratio are shown in wakefulness across rats and 3 recording sessions (W_r1_ W_r2_ W_r3_), during the presence of both behavioral responsiveness and high cortical complexity. **(C, D)** The same putative correlations are shown across rats and across conditions of wakefulness (W) and low ketamine anesthesia (K1), when loss of behavioural responsiveness occurred, but high cortical complexity persisted. **(E, F)** Correlations between PCI^ST^ and HF/LF ratio are also computed and shown across rats and across conditions of low and high ketamine doses (K1 and K2 respectively), when reduction of cortical complexity occurred and behavioral unresponsiveness was unchanged. In panels **A, C, E** the colour maps show the spatial interpolation of R^2^ and *t* scores (a*bove and below respectively, for all panels*) from the correlations of all channels. The horizontal black line in the colour bar indicates the threshold for statistical significance. The white arrowhead indicates channels with statistically significant correlation between PCI^ST^ and HF/LF ratio. In panels **B, D, F** the correlations and/or absence of correlation of right secondary motor cortex (M2_C_, *left side of the panel*) and left posteromedial cortex (RS/V2, *right side of the panel*) are reported with relative R^2^ and *P* values, (corrected for multiple comparisons). Dashed line is used to indicate a correlation close to statistical significance (panel **B**, *left*), while a continuous line indicates a statistically significant linear fitting and correlation (panel **F**, *right*).

In principle, the reduction of HF/LF ratio can be determined by an increase of LF powers, or by a decrease of HF powers or by a combination of the two events. Thus, in order to explain the reduction seen in the posteromedial cortex, we assessed the LF and HF powers at the level of each cortical area, across conditions (**Fig. 5A**). At first, we tested for a possible lateralization, without finding any clear difference between left and right hemispheres in each experimental condition, for both LF powers (**Fig. 5A**; two-way ANOVA, principal effect of lateralization, in W: P = 0.7581, in K1: P = 0.9459, in K2: P = 0.9589;), as well as for HF power (**Fig. 5A**; two-way ANOVA, principal effect of lateralization, in W: P = 0.8284, in K1: P = 0.9664, in K2: P = 0.6748). However, LF powers were differentially distributed across cortical areas, and an overall increase in power could be detected in relation to the increment of ketamine dosage (**Fig. 5B**; two-way rANOVA, principal effect of cortical areas: P = 0.0009, principal effect of ketamine dosage: P = 0.0132). Similar effects were also found for HF powers (**Fig. 5B**; two-way rANOVA, principal effect of cortical areas: P = 0.0109, principal effect of ketamine dosage: P = 0.0041), and were in line with the scaling up of the mean periodograms, seen by averaging across electrodes (**Fig. 1**). Nevertheless, by comparing powers between K1 and K2 conditions for each single channel, a significant increase of LF power was only found at the level of bilateral posteromedial cortex (left and right RS/V2), while the increase of HF power was more spatially sparse, without a clear clusterization (**Fig. 5B**). To more directly compare the relative increment of powers that occurred by increasing ketamine dosage, we computed the ratio between K2 and K1 conditions (K2/K1), for both HF and LF powers, at the level of each cortical area (**Fig. 5C**). Overall, the power increments were differentially distributed across cortical areas, and an overall difference between HF and LF could not be detected (**Fig. 5C**; two-way rANOVA, principal effect of cortical areas: P = 0.0325, principal effect of frequency range: P = 0.0744). However, a clear interaction effect between cortical areas and frequency ranges was identified (**Fig. 5C;** two-way rANOVA, interaction effect: P = 0.0056), thus indicating that the relation between the increments of HF and LF powers changed depending on the specific cortical area. Indeed, LF power increased significantly more than HF power selectively in bilateral posteromedial cortex (left and right RS/V2), conversely HF power had the tendency to increase more than LF power in parieto-frontal areas, even if without reaching statistical significance (**Fig. 5C**).

**Fig. 5.**
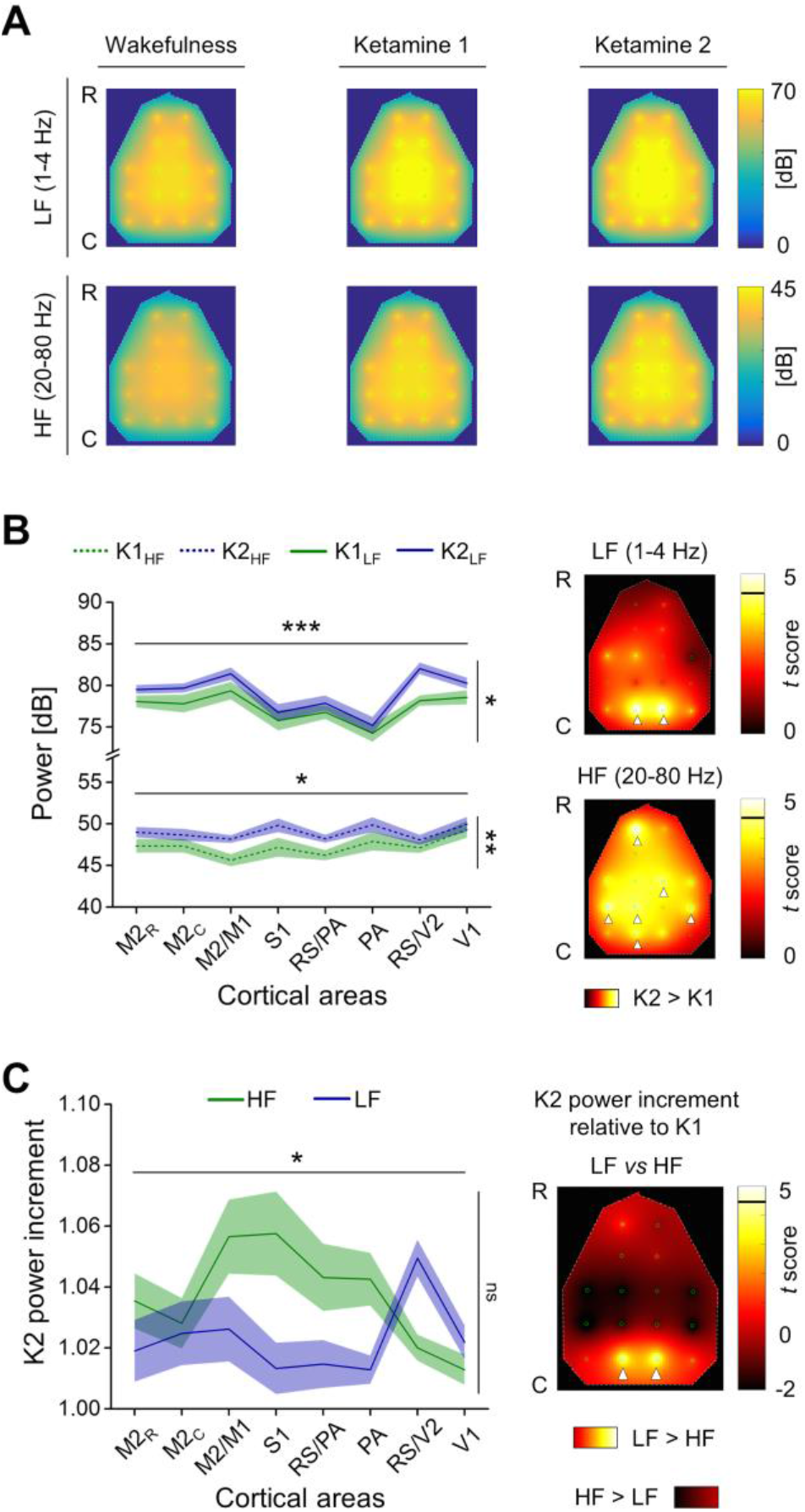
The reduction of HF/LF ratio from low to high dosage of ketamine in the posteromedial cortex was explained by a selective higher increment of LF powers with respect to HF. **(A)** The colour maps show the topographical distributions (R-C: rostral-caudal) of the 16 EEG electrodes (small green circles) and the spatial interpolations of the LF power (1-4 Hz, *above*) and HF power (20-80 Hz, *below*), averaged across time, trials and rats, for each condition. **(B)** *Left*, averaged HF (*dashed line*) and LF (*continuous line*) powers across hemispheres and rats are shown for each cortical area, during both low and high ketamine dosage (K1 and K2 respectively). On the *right*, the colour maps show the spatial interpolation of *t* scores from comparing LF powers (*up*) and HF powers (*bottom*) between K1 and K2 conditions, for each channel. The horizontal black lines in the colour bar indicate the threshold for statistical significance (*t*_*5*_ = 4.5258, Bonferroni-Holm corrected). White arrowheads indicate channels with statistically significant difference across conditions. **(C)** *Left*, the ratio between low and high doses of ketamine is reported for both HF and LF powers at the level of each cortical area, thus showing the power increments induced by deep ketamine anaesthesia (K2). R*ight,* spatial interpolation of the *t* scores from comparing LF power increment with HF power increments induced by increasing ketamine dosage, for each channel. The horizontal black line in the colour bar indicates the threshold for statistical significance (*t*_*5*_ = 4.5258, Bonferroni-Holm corrected). White arrowheads indicate the channels with statistically significant difference between the two frequency ranges.

## DISCUSSION

We reanalysed multichannel EEG data from wakefulness and two levels of ketamine anaesthesia in rats, based on previous experiments (Arena *et al.*, 2020). To assess the capacity for consciousness in each state, we computed PCI^ST^ (Comolatti et al. 2019), and compared these results to region-specific estimates of cortical activation, assessed by HF/LF ratio of spontaneous EEG activity for each electrode (Fernandez *et al.*, 2017; Siclari *et al.*, 2017; Poulet and Crochet, 2018).

In humans, ketamine anaesthesia has been observed to induce a dissociated state of behavioural unresponsiveness with dream-like, vivid experiences (Collier, 1972; Sarasso *et al.*, 2015). We took advantage of this, by using controlled intravenous infusion of ketamine to dissociate the capacity for consciousness from responsiveness. Consistently with human data (Sarasso *et al.*, 2015) we found that PCI^ST^ was not significantly changed relative to wakefulness during the unresponsive state caused by light ketamine anaesthesia (**Fig. 2**), even if it tended to be lower in the K1 condition. This tendency is more in line with results from a larger dataset from rats (Arena *et al.*, 2020), and in agreement with the time course of PCI^ST^, that showed both periods of similarities and differences between wakefulness and light ketamine anaesthesia. The difference in statistical significance of the PCI^ST^ values reported here with respect to previous experiments (Arena *et al.*, 2020) might be due to the smaller sample size here used. However, in both W and K1 conditions, PCI^ST^ build up again after a first decay, reaching similar values between 200 and 300 ms after the stimulus onset (**Fig. 2**). Indeed, our results support the idea that high perturbational complexity is associated with a capacity to sustain long-lasting sequences of deterministic activations (Pigorini *et al.*, 2015; Rosanova *et al.*, 2018), as shown by the durable phase-locked cortical ERPs (**Fig. 2**) and by the strong yet diverse global connectivity observed here **(Supplementary Fig. 2)**. Moreover, the type of response to stimulation required for high PCI^ST^ may also indicate a capacity for the kind of global broadcasting required for consciousness according to GNW (Mashour et al. 2020), as well as for the brain to function as an integrated and differentiated whole as is required for consciousness in IIT (Casali et al. 2013; Tononi et al. 2016). Although a partially reduced level of consciousness might be inferred from the tendency of PCI^ST^ to be lower with light ketamine anaesthesia compared to wakefulness, we observed relatively high spatiotemporal complexity during both conditions, with durable and well integrated phase-locked cortical activations. Thus, taken together, our findings are compatible with a fully, or at least partially, preserved capacity for consciousness during both wakefulness and light ketamine anaesthesia, independently of behavioural responsiveness.

However, ketamine can produce fluctuations of the spatiotemporal complexity of spontaneous EEG in humans, following a bolus injection of an anaesthetic dose (Li and Mashour, 2019), possibly affecting also the capacity for consciousness (Tononi and Edelman, 1998; Ferenets *et al.*, 2006; Schartner *et al.*, 2015). This phenomenon may be caused by unstable pharmacokinetic of the bolus injection, suggesting a dose-dependent effect. We tested this hypothesis by repeating the same electrophysiological recording/stimulations in the same rats during the constant intravenous infusion of ketamine at a higher rate, which gives a more constant systemic concentration. In this high-dose condition, we found that PCI^ST^ strongly reduced along with an earlier interruption of phase-locked response (**Fig. 2**) and with a drastic reduction of cortical functional connectivity and diversity (**Supplementary Fig. 2**). This suggests a reduced capacity to integrate or broadcast information within the global cortical network, and thus a reduced capacity for consciousness during deep ketamine anaesthesia, also indicating a dose-dependent effect.

The spontaneous EEG activity during ketamine anaesthesia was characterized by a widespread, dose-dependent increase in HF power compared to wakefulness (**Fig. 1, 5**). HF oscillations are usually associated with tonic neuronal firing (Steriade, Amzica and Contreras, 1996; Steriade, Timofeev and Grenier, 2001; Mukovski *et al.*, 2007) and cortical activation (Fernandez *et al.*, 2017; Siclari *et al.*, 2017; Poulet and Crochet, 2018). Thus, the observed increase in HF power is consistent with the enhanced presynaptic release occurring after ketamine administration (Ferro *et al.*, 2017), and with the idea that ketamine might mainly inhibit GABAergic interneurons, resulting in a state of overall cortical excitation (Seamans, 2008). Interestingly, LF power also increased from wakefulness to ketamine anaesthesia in a dose dependent manner (**Fig. 1, 5**). Thus, ketamine induced a scaling up of the entire power spectrum of the spontaneous EEG, suggesting a maintained balance between the inhibition and excitation of the overall cortical network underlying the EEG signal (Gao, Peterson and Voytek, 2017). This was supported by the observation of a similar spectral exponent across conditions (**Fig. 1**), which has been hypothesized to indicate an aroused or conscious state (Colombo et al., 2019; Lendner et al., 2020; Arena *et al.*, 2020). Why then did we observe the strong reduction of PCI^ST^ in the transition from low to high ketamine condition (**Fig. 2**)?

One hypothesis is that specific cortical circuits might be particularly relevant for sustaining complex neuronal interactions, and that the activation state of these circuits might diverge from the average dynamic of the entire cortical network. Indeed, it is known that transient and local cortical deactivations or activations can occur and dissociate from the global brain state, such as with local sleep during wakefulness (Murphy *et al.*, 2011; Vyazovskiy *et al.*, 2011; Fernandez *et al.*, 2017), possibly modifying the capacity for behaviour and/or conscious experience (Vyazovskiy *et al.*, 2011; Fernandez *et al.*, 2017; Siclari *et al.*, 2017; Poulet and Crochet, 2018). For example, it has been shown that localized reduction of LF power and increased HF activity (high HF/LF ratio) within the posterior cortex is strongly associated with dream experience in humans (Siclari *et al.*, 2017) during NREM sleep, a state dominated by LF activity that is often linked to unconsciousness (Tononi and Massimini, 2008). Thus, to uncover the role and the state of activation of specific areas, we similarly measured the HF (20-80 Hz) / LF (1-4 Hz) power ratio from the spontaneous activity of all the 16 epidural electrodes, and compared across experimental conditions.

Although both the light and deep ketamine anesthesia caused an unresponsive behavioral state, we were able to identify regional variations in HF/LF ratio that were related to changes in the global PCI^ST^ value. Strikingly, the bilateral posteromedial cortex was the only region that showed a consistent reduction of HF/LF ratio, from low to high ketamine dosage, along with the drop in PCI^ST^ (**Fig. 3**). The reduced ratio within the posteromedial cortex indicated a local deactivation (Poulet and Crochet, 2018), which was explained by a larger increase of LF than HF power induced by the increased ketamine dosage (**Fig. 5**). Consistently, the HF/LF ratio over the posteromedial cortex strongly correlated with the PCI^ST^ level during ketamine administration (**Fig. 4**) and across all conditions (**Fig. 3)**, but not within wakefulness alone or between wakefulness and light ketamine anesthesia (**Fig. 4**), when PCI^ST^ did not change substantially. These results indicate that the state of the posteromedial cortex is strongy correated with the capacity for long-lasting, complex cortical activations and thus possibly, according to IIT, may play an important role in sustaining the capacity for consciousness. In other words, our results may support the hypothesis that a selective deactivation of the posteromedial cortex - as indicated by the localized decrease in HF/LF power - is correlated with, and may even underlie, a sharp reduction of the brain’s capacity to function as an integrated whole that is capable of sustaining consciousness. In contrast, light ketamine anaesthesia produced a significant reduction of HF/LF ratio compared to wakefulness only over the right primary motor and somatosensory cortex (**Fig. 3**). This was consistent with the loss of behavioural responsiveness induced by the low ketamine dose, and possibly with an analgesic effect. A correlation between the HF/LF ratio of the right secondary motor cortex and PCI^ST^ was also found across conditions (**Fig. 3**). However, this relation was at least partially explained by a wakefulness-specific weak correlation (**Fig. 4**), which could reflect variations within the same state, such as active/quite wakefulness or transients attentional loading.

In the current experimental setting, the electrodes over the posteromedial cortex cover both the medial part of secondary visual cortex and the caudal part of retrosplenial cortex (**Supplementary Fig. 1**). Interestingly, the dose-dependent deactivation of this cortical region is reminiscent of a recent finding, in which sub-anaesthetic bolus injections of ketamine were found to induce slow wave oscillations and synchronized, rhythmic neuronal silences selectively in retrosplenial cortex in mice, affecting behaviour (Vesuna *et al.*, 2020). Besides, the retrosplenial cortex is a particularly highly integrated area, within the medial cortical subnetwork in the rodent brain (Zingg *et al.*, 2014). It receives information from claustrum, indirectly from the hippocampus through subiculum, and it is directly interconnected with several sensory areas (visual, auditory and somatosensory) and high-order associative areas, including medial frontal cortex. Thus, it is likely to play important roles in multisensory integration with also higher functions, such as episodic memory, spatial navigation and motor planning (Zingg *et al.*, 2014). Given this, it is not surprising that a deactivation of this area (low HF/LF ratio, **Fig. 3**), due to enhanced LF activity (**Fig. 5**), could go with the disruption of widespread integration of complex cortical interactions as seen here, with the drop of PCI^ST^ at high ketamine dosage (**Fig. 2**). In other words, there is reason to believe that the specific deactivation of a region in the posteromedial cortex can be directly involved in breaking down the properties required for sustaining a capacity for consciousness.

The main findings presented here are compatible with several theories of consciousness, as it is widely agreed that some sort of long range interactions within the brain are required to sustain its capacity for consciousness. For example, GNW requires information to be globally broadcast (Mashour *et al.*, 2020), IIT requires the physical substrate of consciousness to be integrated (Tononi *et al.*, 2016), and at least some higher order theories require long-range interaction to maintain the capacity to form representations in associative cortices about first order states in early sensory regions (Lau and Rosenthal, 2011, Brown et al., 2019). Our results suggest that the kind of global integration typically associated with vivid conscious experience, breaks down when ketamine specifically deactivates the posteromedial cortex (**Fig. 3, 4**). Thus, any theory that considers PCI^ST^ as a reliable marker of the capacity for consciousness must grapple with the finding that high doses of ketamine appears to lead to a loss of consciousness associated with a specific deactivation of the posteromedial cortex.

While the claim that the integrity of the posterior cortex is necessary for the capacity for consciousness may not by itself distinguish between theories, other findings reported here may be more decisive. Specifically, the finding that the ketamine-induced deactivation of primary motor and somatosensory cortex was associated with loss of behavioural responsiveness, but no significant change in PCI^ST^ (see **Fig. 3**), indicates that some cortices may be deactivated without disrupting the normal capacity for consciousness. Thus, to be consistent with our data, theories of consciousness should also be able to explain why some regions may be deactivated without any apparent effect on the brain’s overall capacity for consciousness. Furthermore, we observed that changes in the HF/LF ratio of secondary motor regions during wakefulness weakly correlated with changes in PCI^ST^ (**Fig. 4**). This relation may reflect a modulatory effect of frontal regions on the overall capacity for consciousness during wakefulness. Even though these variations were not the results of controlled intervention, they suggest that spontaneous activity (HF/LF) changes in the front of the cortex, with its associated cognitive functions involving working memory, cognitive control, planning and decision-making, attention, etc. (Dalley et al. 2004), might modulate cortical complexity within limits of what we consider normal capacity for consciousness during wakefulness.

Of course, these findings are not conclusive, as the observed minor changes in HF/LF may only suggest regional fluctuations as opposed to interventional inactivation. Furthermore, it is not necessarily the case that the changes in HF/LF observed in different regions were caused by the same underlying processes, and it is also uncertain that they were always indicative of a deactivation of the region. To alleviate these issues, we aim to perform experiments in the future with controlled, direct inactivation of individual cortical regions while measuring PCI^ST^ from awake rodents. While awaiting the results, we invite proponents of theories of consciousness to respond to the following challenge: what effect does your favourite theory predict that controlled deactivations of specific regions of cortex will have on a rodent’s overall capacity for consciousness, as measured by PCI^ST^? Specifically, which regions will, when inactivated, lead to a strong reduction in the global PCI^ST^, which will have only a smaller or modulatory effect, and which will have no significant effect on the global PCI^ST^?

## CONCLUSIONS

By comparing EEG in light and deep ketamine anaesthesia in rats, we dissociated changes in global and local cortical dynamics related to loss of behavioural responsiveness from those likely related to consciousness (assessed by PCI^ST^). Light ketamine anaesthesia induced an unresponsive state with apparent deactivation of primary somatosensory and motor regions (indicated by locally reduced HF/LF power ratio). The mean PCI^ST^ value was somewhat lower than in wakefulness, but long-lasting phase-locked responses, with high and diversified functional connectivity suggested a dissociated state, possibly with only a partially reduced consciousness level. In contrast, deep ketamine anaesthesia strongly reduced PCI^ST^, with early interruption of phase-locked response and disruption of functional connectivity, suggesting a more fully unconscious state. This was associated with a highly selective deactivation of the posteromedial cortex, compared to light ketamine anaesthesia, suggesting a primary role for this cortical region in the capacity for consciousness. Thus, we found evidence for regionally specific effects of deactivating cortical areas on the capacity for consciousness, which theories of consciousness should be able to account for.

## ADDITIONAL INFORMATION

### Authors’ Contribution

AA designed the experiments; AA, ST performed experiments and collected data; AA analyzed data; RC performed PCIST analysis; AA, BEJ, JFS, wrote the manuscript; all authors participated in the interpretation of results and revision of the manuscript, and approved the final version of the manuscript.

### Funding

This project/research received funding from the European Union’s Horizon 2020 Framework Programme for Research and Innovation under the Specific Grant Agreements No. 785907 (Human Brain Project SGA2; JFS), No. 720270 (Human Brain Project SGA1; JFS)

### Competing interests

The authors declare that they have no financial competing interests.

### Data availability

Data will be available in the public repository EBRAINS upon publication.

## Notes

### Competing Interest Statement

The authors have declared no competing interest.

## Bibliography

Arena, A. et al. (2020) ‘General anaesthesia disrupts complex cortical dynamics in response to intracranial electrical stimulation in rats’, biorxiv - submitted and under peer review. doi: 10.1101/2020.02.25.964056.

Arena, A., Thon, S. and Storm, J. (2019) ‘PCI-like measure in rodents’. Human Brain Project Neuroinformatics Platform. doi: 10.25493/S0DM-BK5.

Blumenfeld, H. (2005) ‘Consciousness and epilepsy: why are patients with absence seizures absent?’, Progress in brain research, 150, pp. 271–286.

Boly, M. et al. (2017) ‘Are the Neural Correlates of Consciousness in the Front or in the Back of the Cerebral Cortex? Clinical and Neuroimaging Evidence’, The Journal of neuroscience: the official journal of the Society for Neuroscience, 37(40), pp. 9603–9613.

Brown, E. N., Lydic, R. and Schiff, N. D. (2010) ‘General anesthesia, sleep, and coma’, The New England journal of medicine, 363(27), pp. 2638–2650.

Brown, R., Lau, H. and LeDoux, J. E. (2019) ‘Understanding the Higher-Order Approach to Consciousness’, Trends in cognitive sciences, 23(9), pp. 754–768.

Casali, A. G. et al. (2013) ‘A theoretically based index of consciousness independent of sensory processing and behavior’, Science translational medicine, 5(198), p. 198ra105.

Casarotto, S. et al. (2016) ‘Stratification of unresponsive patients by an independently validated index of brain complexity’, Annals of neurology, 80(5), pp. 718–729.

Chernik, D. A. et al. (1990) ‘Validity and reliability of the Observer’s Assessment of Alertness/Sedation Scale: study with intravenous midazolam’, Journal of clinical psychopharmacology, 10(4), pp. 244–251.

Cohen, M. X. (2014) Analyzing Neural Time Series Data: Theory and Practice. 1 edition. Cambridge, Massachusetts: MIT Press.

Collier, B. B. (1972) ‘Ketamine and the conscious mind’, Anaesthesia, 27(2), pp. 120–134.

Colombo, M. A. et al. (2019) ‘The spectral exponent of the resting EEG indexes the presence of consciousness during unresponsiveness induced by propofol, xenon, and ketamine’, NeuroImage, 189, pp. 631–644.

Comolatti, R. et al. (2019) ‘A fast and general method to empirically estimate the complexity of brain responses to transcranial and intracranial stimulations’, Brain stimulation, 12(5), pp. 1280–1289.

Compte, A. et al. (2003) ‘Cellular and network mechanisms of slow oscillatory activity (<1 Hz) and wave propagations in a cortical network model’, Journal of neurophysiology, 89(5), pp. 2707–2725.

David, O., Kilner, J. M. and Friston, K. J. (2006) ‘Mechanisms of evoked and induced responses in MEG/EEG’, NeuroImage, 31(4), pp. 1580–1591.

Del Cul, A. et al. (2009) ‘Causal role of prefrontal cortex in the threshold for access to consciousness’, Brain: a journal of neurology, 132(Pt 9), pp. 2531–2540.

Esser, S. K., Hill, S. L. and Tononi, G. (2007) ‘Sleep homeostasis and cortical synchronization: I. Modeling the effects of synaptic strength on sleep slow waves’, Sleep, 30(12), pp. 1617–1630.

Ferenets, R. et al. (2006) ‘Comparison of entropy and complexity measures for the assessment of depth of sedation’, IEEE transactions on bio-medical engineering, 53(6), pp. 1067–1077.

Fernandez, L. M. J. et al. (2017) ‘Highly Dynamic Spatiotemporal Organization of Low-Frequency Activities During Behavioral States in the Mouse Cerebral Cortex’, Cerebral cortex, 27(12), pp. 5444–5462.

Ferro, M. et al. (2017) ‘Functional mapping of brain synapses by the enriching activity-marker SynaptoZip’, Nature communications, 8(1), p. 1229.

Funk, C. M. et al. (2017) ‘Role of Somatostatin-Positive Cortical Interneurons in the Generation of Sleep Slow Waves’, The Journal of neuroscience: the official journal of the Society for Neuroscience, 37(38), pp. 9132–9148.

Gao, R., Peterson, E. J. and Voytek, B. (2017) ‘Inferring synaptic excitation/inhibition balance from field potentials’, NeuroImage, 158, pp. 70–78.

Gao, S. and Calderon, D. P. (2020) ‘Robust alternative to the righting reflex to assess arousal in rodents’, Scientific reports, 10(1), p. 20280.

Ghoneim, M. M. et al. (2009) ‘Awareness during anesthesia: risk factors, causes and sequelae: a review of reported cases in the literature’, Anesthesia and analgesia, 108(2), pp. 527–535.

Giacino, J. T., Kalmar, K. and Whyte, J. (2004) ‘The JFK Coma Recovery Scale-Revised: Measurement characteristics and diagnostic utility’, Archives of physical medicine and rehabilitation, 85(12), pp. 2020–2029.

Koch, C. et al. (2016) ‘Neural correlates of consciousness: progress and problems’, Nature reviews. Neuroscience, 17(5), pp. 307–321.

Lau, H. and Rosenthal, D. (2011) ‘Empirical support for higher-order theories of conscious awareness’, Trends in cognitive sciences, 15(8), pp. 365–373.

Laureys, S. et al. (2010) ‘Unresponsive wakefulness syndrome: a new name for the vegetative state or apallic syndrome’, BMC medicine, 8, p. 68.

Lendner, J. D. et al. (2020) ‘An electrophysiological marker of arousal level in humans’, eLife, 9. doi: 10.7554/eLife.55092.

Li, D. and Mashour, G. A. (2019) ‘Cortical dynamics during psychedelic and anesthetized states induced by ketamine’, NeuroImage, 196, pp. 32–40.

Loomis, A. L., Harvey, E. N. and Hobart, G. A. (1937) ‘Cerebral states during sleep, as studied by human brain potentials’, Journal of experimental psychology, 21(2), p. 127.

Mashour, G. A. et al. (2020) ‘Conscious Processing and the Global Neuronal Workspace Hypothesis’, Neuron, 105(5), pp. 776–798.

Massimini, M. et al. (2004) ‘The sleep slow oscillation as a traveling wave’, The Journal of neuroscience: the official journal of the Society for Neuroscience, 24(31), pp. 6862–6870.

Massimini, M. et al. (2009) ‘A perturbational approach for evaluating the brain’s capacity for consciousness’, Progress in brain research, 177, pp. 201–214.

Mukovski, M. et al. (2007) ‘Detection of active and silent states in neocortical neurons from the field potential signal during slow-wave sleep’, Cerebral cortex, 17(2), pp. 400–414.

Murphy, M. et al. (2011) ‘The cortical topography of local sleep’, Current topics in medicinal chemistry, 11(19), pp. 2438–2446.

Nielsen, T. A. (2000) ‘A review of mentation in REM and NREM sleep: “covert” REM sleep as a possible reconciliation of two opposing models’, The Behavioral and brain sciences, 23(6), pp. 851–66; discussion 904–1121.

Noreika, V. et al. (2011) ‘Consciousness lost and found: subjective experiences in an unresponsive state’, Brain and cognition, 77(3), pp. 327–334.

Odegaard, B., Knight, R. T. and Lau, H. (2017) ‘Should a Few Null Findings Falsify Prefrontal Theories of Conscious Perception?’, The Journal of neuroscience: the official journal of the Society for Neuroscience, 37(40), pp. 9593–9602.

Paxinos, G. and Watson, C. (2007) The Rat Brain in Stereotaxic Coordinates. 6th Edition. Academic Press.

Pigorini, A. et al. (2015) ‘Bistability breaks-off deterministic responses to intracortical stimulation during non-REM sleep’, NeuroImage, 112, pp. 105–113.

Poulet, J. F. A. and Crochet, S. (2018) ‘The Cortical States of Wakefulness’, Frontiers in systems neuroscience, 12, p. 64.

Rosanova, M. et al. (2018) ‘Sleep-like cortical OFF-periods disrupt causality and complexity in the brain of unresponsive wakefulness syndrome patients’, Nature communications, 9(1), p. 4427.

Sanders, R. D. et al. (2012) ‘Unresponsiveness ≠ unconsciousness’, Anesthesiology, 116(4), pp. 946–959.

Sarasso, S. et al. (2015) ‘Consciousness and Complexity during Unresponsiveness Induced by Propofol, Xenon, and Ketamine’, Current biology: CB, 25(23), pp. 3099–3105.

Schartner, M. M. et al. (2015) ‘Complexity of Multi-Dimensional Spontaneous EEG Decreases during Propofol Induced General Anaesthesia’, PloS one, 10(8), p. e0133532.

Seamans, J. (2008) ‘Losing inhibition with ketamine’, Nature chemical biology, 4(2), pp. 91–93.

Siclari, F. et al. (2017) ‘The neural correlates of dreaming’, Nature neuroscience, 20(6), pp. 872–878.

Siegle, J. H. et al. (2017) ‘Open Ephys: an open-source, plugin-based platform for multichannel electrophysiology’, Journal of neural engineering, 14(4), p. 045003.

Steriade, M., Amzica, F. and Contreras, D. (1996) ‘Synchronization of fast (30-40 Hz) spontaneous cortical rhythms during brain activation’, The Journal of neuroscience: the official journal of the Society for Neuroscience, 16(1), pp. 392–417.

Steriade, M., Nuñez, A. and Amzica, F. (1993) ‘A novel slow (< 1 Hz) oscillation of neocortical neurons in vivo: depolarizing and hyperpolarizing components’, The Journal of neuroscience: the official journal of the Society for Neuroscience, 13(8), pp. 3252–3265.

Steriade, M., Timofeev, I. and Grenier, F. (2001) ‘Natural waking and sleep states: a view from inside neocortical neurons’, Journal of neurophysiology, 85(5), pp. 1969–1985.

Tononi, G. (2004) ‘An information integration theory of consciousness’, BMC neuroscience, 5(1), p. 42.

Tononi, G. et al. (2016) ‘Integrated information theory: from consciousness to its physical substrate’, Nature reviews. Neuroscience, 17(7), pp. 450–461.

Tononi, G. and Edelman, G. M. (1998) ‘Consciousness and complexity’, Science, 282(5395), pp. 1846–1851.

Tononi, G. and Massimini, M. (2008) ‘Why does consciousness fade in early sleep?’, Annals of the New York Academy of Sciences, 1129, pp. 330–334.

Vanhaudenhuyse, A. et al. (2018) ‘Conscious While Being Considered in an Unresponsive Wakefulness Syndrome for 20 Years’, Frontiers in neurology, 9, p. 671.

Vesuna, S. et al. (2020) ‘Deep posteromedial cortical rhythm in dissociation’, Nature, 586(7827), pp. 87–94.

Volgushev, M. et al. (2006) ‘Precise Long-Range Synchronization of Activity and Silence in Neocortical Neurons during Slow-Wave Sleep’, The Journal of neuroscience: the official journal of the Society for Neuroscience, 26(21), pp. 5665–5672.

Vyazovskiy, V. V. et al. (2009) ‘Cortical firing and sleep homeostasis’, Neuron, 63(6), pp. 865–878.

Vyazovskiy, V. V. et al. (2011) ‘Local sleep in awake rats’, Nature, 472(7344), pp. 443–447.

Zingg, B. et al. (2014) ‘Neural networks of the mouse neocortex’, Cell, 156(5), pp. 1096–1111.

