## Supplementary Fig. for "Does the posteromedial cortex play a primary role for the capacity for consciousness in rats?"

### SUPPLEMENTARY FIGURES:

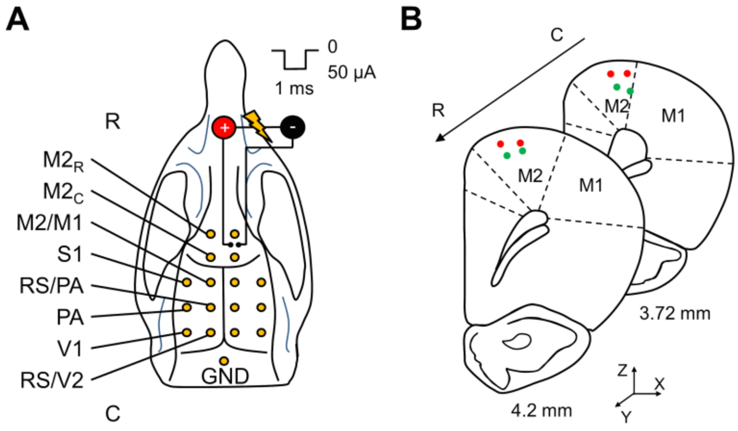


**Supplementary Fig. 1 Location of recording and stimulating electrodes (A)** Positions in the rat skull of the 16 screw electrodes (yellow dots) and bipolar stimulating electrode used for recording EEG and triggering ERPs (R-C: rostral-caudal). Recording electrodes were in contact with the dura and were organized in a grid, symmetric along the sagittal suture, covering most of the cortical surface. Recording electrodes were placed at the following coordinates with respect to Bregma (in mm; X is the medio-lateral axis and Y is the rostro-caudal axis ): X = ± 1.5, Y = + 5 (rostral part of secondary motor cortex, $M2_R$); X = ± 1.5, Y = + 2 (caudal part of secondary motor cortex, $M2_C$); X = ± 1.5, Y = - 1 (secondary and primary motor cortex, M2/M1); X = ± 4.5, Y = - 1 (primary somatosensory cortex, S1); X = ± 1.5, Y = - 4 (retrosplenial and parietal associative cortex, RS/PA); X = ± 4.5, Y = - 4 (parietal associative cortex, PA); X = ± 1.5, Y = - 7 (retrosplenial and secondary visual cortex, V2); X = ± 4.5, Y = - 7 (primary visual cortex, V1); X = 0, Y = - 10 (ground, GND, over cerebellum). The bipolar tungsten electrode used for stimulation (electrical pulses of 50 µA, 1 ms, triggered at 0.1 Hz) was inserted perpendicularly to the cortical surface, along the coronal plane in the right secondary motor cortex (M2), at the coordinates (in mm from Bregma; Z is the dorso-ventral axis): X = + 1.2 left wire / + 1.7 right wire; Y = + 3.7; Z = + 1.9. **(B)** Representation of coronal brain sections of the rat brain (one hemisphere) with actual position of bipolar electrodes of 4 rats (dots represent the 2 poles of the bipolar electrode and each rat is colour coded in each slice). Coordinates have been measured from Nissl stained coronal sections of rat brains.

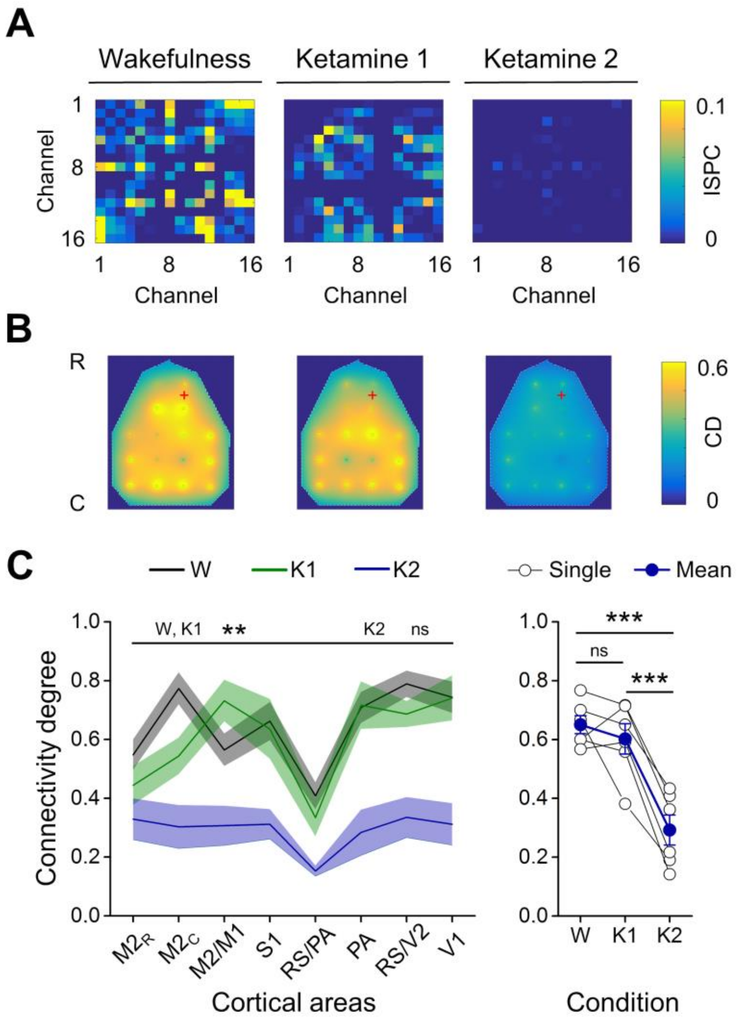


**Supplementary Fig. 2 The degree of connectivity and diversity in response to stimulation was similar between wakefulness and low dosage of ketamine, but collapsed by increasing ketamine dosage. (A)** The functional connectivity matrices of one rat during wakefulness and with low and high doses of ketamine are reported. The channel-by-channel matrices show the inter-site phase clustering (ISPC) across ERPs for each channel pair, averaged in frequency range 5-14 Hz and in time window 180-400 ms. The time range extends beyond the end of the HF suppression after stimulation (172.809 ± 20.777 ms, mean across rats ± SEM), up to the drop of the phase locking across trials (369.762 ± 59.859 ms, mean across rats ± SEM) in light ketamine anaesthesia. It also included the late increase of HF power for all conditions (mean onset across rats and conditions: 212.03 ± 21.17 ms). **(B)** The topographical distribution (R-C: rostral-caudal) of the connectivity degree (CD) averaged across rats for each channel is interpolated and shown for each condition of **(A)**. The red cross indicates the stimulation site. For each channel, the connectivity degree indicates the proportion of significant synchronized channel pairs (mean ISPC > 0). **(C)** *Left***,** the average CD across hemispheres and rats are shown for each cortical area and condition (shade is SEM across rats).

During both wakefulness and light ketamine anaesthesia, the distribution of CD is differentiated across cortical areas while is more homogeneous with high dosage of ketamine (6 rats; one-way rANOVA, principal effect of cortical areas, in W: $P$ = 0.002, in K1: $P$ = 0.0018 in K2: $P$ = 0.1639). *Right,* The average CDs across channels are reported for all rats and conditions and show a similar degree of functional connectivity between wakefulness and light ketamine anaesthesia, that is strongly reduced by increasing the dosage of ketamine (6 rats; one-way rANOVA, principal effect of state condition, $P = 1.9548*10^{-5}$; paired samples t-test, W *vs* K1, $P$ = 0.4165, W *vs* K2, $P$ = 0.0008, K1 *vs* K2, $P$ = 0.0008).

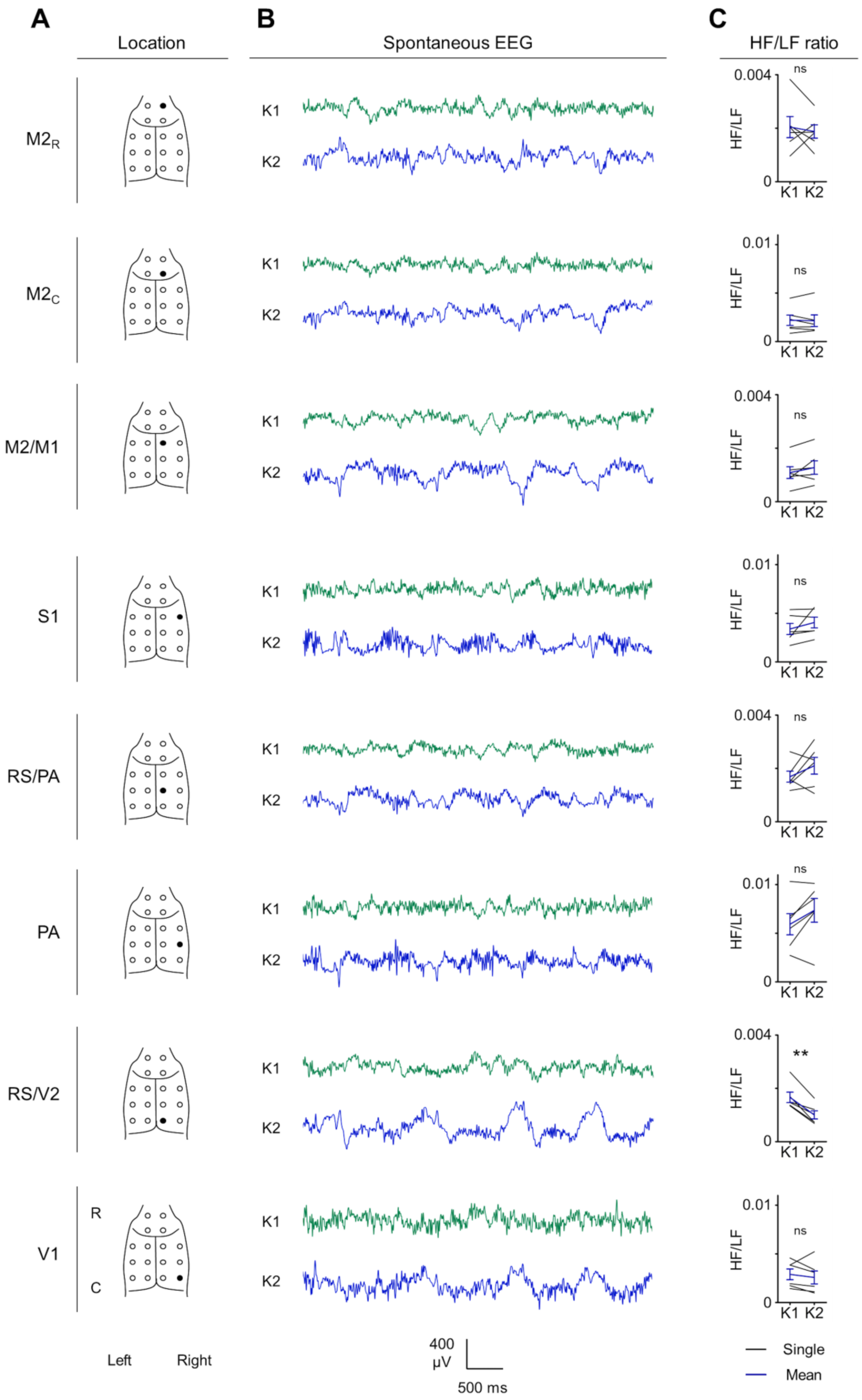
A
Location
B
Spontaneous EEG
C
HF/LF ratio
M2R
M2C
M2/M1
S1
RS/PA
PA
RS/V2
V1
K1
K2
0.004
0.01
0
ns
**
HF/LF
R
C
Left
Right
400 µV
500 ms
Single
Mean

**Supplementary Fig. 3 Spontaneous EEG and HF/LF ratio from each cortical area at incremental doses of ketamine (A)** Representation of the 16 EEG electrodes (dots) and their positions over the rat skull (R-C: rostral-caudal). For each row, the black dot indicates the electrode from which the electrophysiological recordings on the right are from. All the electrodes form the right hemisphere are shown (from *up* to *bottom*) and on the left, the names of the corresponding recorded cortical areas are indicated. **(B)** Example of spontaneous EEG (5 seconds) from each electrode of the right hemisphere of one rat during light ketamine anaesthesia (K1) and deep ketamine anaesthesia (K2). **(C)** On the right, the ratio between high frequency (HF, 20-80 Hz) and low frequency (LF, 1-4 Hz) powers (HF/LF ratio), averaged across hemispheres, is shown for all 6 rats and cortical areas (rows). The only cortical area that showed a statistically significant reduction of HF/LF ratio from K1 to K2, was the posteromedial cortex or retrosplenial/secondary visual cortex (RS/V2). The same effect was detected in RS/V2 of both hemispheres and was consistent across the tested rats.

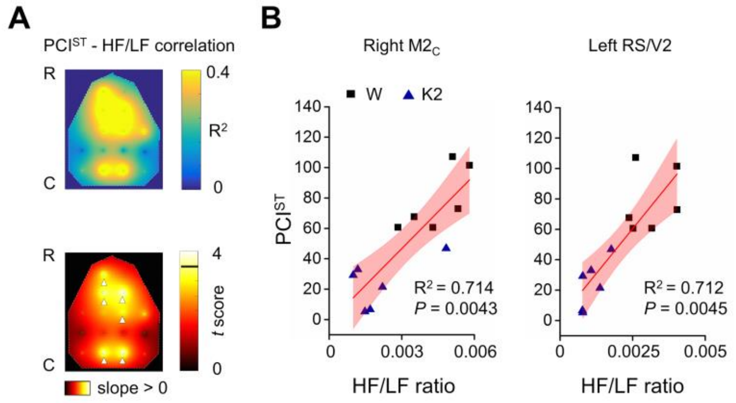


**Supplementary Fig. 4 Both secondary motor cortex and posteromedial cortex showed a correlation between PCI$^{ST}$ and HF/LF ratio when both loss of behavioral responsiveness and loss of cortical complexity occurred. (A)** A*bove,* the colour map shows the spatial distribution of the coefficient of determination $R^2$ from the correlation between PCI$^{ST}$ and HF/LF ratio, for each channel, across rats and conditions of wakefulness (W) and high dosage of ketamine (K2). *Below*, the colour map reports the spatial interpolation of *t* scores (regression t-test) to test $H_0$: slope of linear regression = 0, thus assessing the statistical significance of the correlation for each channel. The horizontal black line in the colour bar indicates the threshold for statistical significance ($t_{10}$ = 3.4477, Bonferroni-Holm corrected). White arrowheads indicate the channels showing a statistically significant correlation between PCI$^{ST}$ and HF/LF ratio. We found that when conditions that differ for both behavioural responsiveness and cortical complexity are considered together (such as W and K2), the HF/LF ratio from the spontaneous EEG of both secondary motor cortex and posteromedial cortex similarly correlated with the level of perturbational complexity of the entire network. The correlations of right $M2_C$ and left RS/V2 are reported in **(B)** with respective $R^2$ and *P* values.
